# Longitudinal Proteomics Analysis of Cerebrospinal Fluid in Survivors of Childhood Acute Lymphoblastic Leukemia

**DOI:** 10.64898/2026.01.22.701141

**Authors:** Mingming Niu, Him K. Shrestha, Hong Wang, Xusheng Wang, Kevin Krull, Junmin Peng

**Author notes:** Co-Corresponding authors: Him K. Shrestha, Kevin Krull, and Junmin Peng.

## Abstract

Mass spectrometry-based proteomic profiling of cerebrospinal fluid (CSF) in large patient cohorts, particularly among childhood cancer survivors, remains scarce, limiting opportunities for benchmarking, method development, and validation. Here, we present a longitudinal CSF proteomics dataset from survivors of childhood acute lymphoblastic leukemia (ALL). Samples were collected from 178 patients enrolled in the TOTXVI therapeutic protocol at two timepoints: diagnosis (pre-treatment) and during the consolidation phase of chemotherapy (totaling 356 samples). CSF proteomes were profiled using tandem-mass-tag (TMT) labeling coupled with extensive fractionation and high-resolution liquid chromatography-tandem mass spectrometry (LC/LC-MS/MS). The dataset includes quantitative profiles for more than 3,000 confidently identified unique proteins. This resource enables the investigation of chemotherapy-induced alterations in the CSF proteome and provides a valuable resource for studying the molecular mechanisms of neurotoxicity, identifying biomarkers of adverse late effects, and guiding the development of neuroprotective strategies in childhood cancer survivors.

## Background & Summary

Childhood acute lymphoblastic leukemia (ALL) is the most common pediatric cancer, and modern chemotherapy has raised survival to nearly 95 percent^1,2^. However, 30-40 percent of survivors develop long-term neurocognitive difficulties, including problems with attention, processing speed, and executive function^3-6^. Preserving cognitive function in ALL survivors is, therefore, as critical as achieving remission, given its profound impact on overall quality of life. These deficits are associated with structural and functional brain abnormalities and are thought to arise, in part, from neurotoxic effects of chemotherapy agents such as methotrexate^6,7^.

Understanding the biological mechanisms underlying these neurocognitive impairments in pediatric ALL remains a critical clinical and scientific challenge. A key step forward requires moving beyond descriptive associations to establish mechanistic insight, specifically by investigating biological pathways and identifying key drivers that drive neuronal damage. Systemic biomarkers, such as those from peripheral blood, may often lack specificity for central nervous system injury^8^. In contrast, cerebrospinal fluid (CSF) provides a uniquely accessible and highly informative tissue compartment in pediatric ALL, as routine lumbar punctures are already integrated into clinical care. Prior targeted studies have shown that CSF markers such as tau, myelin basic protein, and glial fibrillary acidic protein correlate with neurocognitive outcomes and neuroimaging abnormalities, underscoring its value for identifying early indicators of neurotoxicity^9-12^. This highlights the need for deeper CSF proteomic characterization to move beyond selected markers and enable comprehensive mapping of the molecular pathways involved in chemotherapy-induced injury.

Thus, the primary focus of this data resource was to address this gap by providing a comprehensive, unbiased proteomic profile of CSF from childhood ALL patients treated on the TOTXVI therapeutic protocol. The dataset contains longitudinal proteomic data from 178 patients at diagnosis (baseline) and during the consolidation phase of therapy. These data offer a valuable resource for investigating the biological processes of chemotherapy-induced neurotoxicity. The reuse potential includes, but is not limited to, identifying novel biomarkers of central nervous system injury, exploring mechanisms of neurotoxicity, and identifying potential targets for future neuroprotective interventions.

## Methods

### Mass Spectrometry Protein Profiling

CSF samples were collected from patients enrolled in the Total Therapy Study XVI (TOTXVI), a clinical treatment protocol at St. Jude Children’s Research Hospital for pediatric acute lymphoblastic leukemia^13^. In total, 356 CSF samples, including 178 matched pairs, were included in this proteomics study. Each sample was mixed with 8 M urea and vortexed to ensure complete solubilization. This was followed by reduction and alkylation as described previously^14^. Proteins were digested in a two-step enzymatic workflow. First, Lys-C was added at a 1:100 enzyme-to-substrate ratio and incubated under denaturing conditions. Samples were then diluted to reduce the urea concentration below 2 M, and sequencing-grade trypsin was added at a 1:50 enzyme-to-substrate ratio. Samples were incubated overnight at 37 °C to ensure complete digestion. Resulting peptides were desalted using standard reversed-phase C18 solid-phase extraction. Desalted peptides were dried by vacuum centrifugation and stored at -20 °C.

Following desalting, samples were grouped for tandem mass tag (TMT) labeling based on block randomization. A total of 24 batches were prepared for TMTpro labeling, each containing 15 samples and one internal reference for batch normalization. The internal reference was a mixture of a small fraction of representative samples. Briefly, peptides were reconstituted in labeling buffer, reacted with TMTpro reagents according to manufacturer protocols, quenched, and then pooled within each batch. For each TMT batch, pooled peptides were subjected to high-pH reversed-phase chromatography on a 3.0 × 150 mm, 1.7-micrometer C18 column (serial number #03653017616612) over a 120-minute gradient. Peptides were separated into 40 fractions to improve proteome depth. Fractions were dried and reconstituted in 0.1 percent formic acid prior to LC-MS/MS analysis.

Each fraction was analyzed by low pH nanoflow liquid chromatography coupled with high-resolution mass spectrometer (Q Exactive HF). The mass spectrometer is operated in data-dependent mode with a survey scan in Orbitrap (450–1600 m/z, 60,000 resolution, 1 × 10^6^ automatic gain control (AGC), ∼50 ms maximal ion time) and perform 20 data-dependent MS/MS high-resolution scans (60,000 resolution, 1 × 10^5^ AGC target, ∼150 ms maximal ion time, 32% HCD normalized collision energy (NCE), 1.0 m/z isolation window, 0.2 m/z isolation offset, and 10 s dynamic exclusion).

### Peptide and Protein Identification

MS data were analyzed using the JUMP software suite, which is optimized for TMT-based proteomic workflows and supports large-scale searches through high-performance computing clusters^15^. Briefly, raw MS files were converted to mzXML format, and precursor ions were preprocessed to refine charge-state assignments. JUMP then generated peptide tags from MS/MS spectra and performed database searching by matching experimental spectra against theoretical fragmentation patterns, producing ranked peptide-spectrum matches (PSMs) using the Jscore metric. Searches were conducted against human proteome database (UniProt) with a precursor mass tolerance of 10 ppm and a product-ion tolerance of 15 ppm. Additional search parameters included fully tryptic digestion with up to two missed cleavages, a maximum of three modification sites per peptide, and annotation of a, b, and y fragment ions. Static modifications included TMT16 labeling of Lys residues and peptide N termini (+304.20715 Da) and carbamidomethylation of Cys (+57.02146 Da). Methionine oxidation (+15.99491 Da) was included as a variable modification.

PSMs were filtered using JUMP-specific matching scores (Jscore and ΔJn) together with precursor mass accuracy to achieve a protein-level false discovery rate (FDR) <1%. PSMs were further grouped by peptide length (minimum 7 amino acids), precursor charge state, tryptic termini, modifications, and missed cleavages, then re-filtered to maintain an overall FDR of approximately 1 percent. FDR was estimated using a target-decoy approach^16,17^. For peptides shared across multiple proteins, the protein with the highest number of supporting PSMs was selected following the principle of parsimony. Peptide and protein quantification were obtained by averaging TMT reporter-ion intensities across all matched PSMs.

### Protein quantification, data imputation, and covariate correction

Protein quantification values were exported as Excel tables and log_2_-transformed, followed by normalization using an internal reference. Proteins missing in >50 percent of samples were removed, protein isoforms were consolidated, and remaining missing values were imputed using K-nearest neighbors (KNN) implemented via sklearn.impute in Python’s scikit-learn package^18^. After imputation, protein abundances were mean centered across samples. Covariates such as age, sex, and batch were corrected using a linear regression model^19^. Differential abundance analysis was performed using moderated paired t-tests comparing pre- and post-treatment samples, implemented with the *limma* package in RStudio^20^. Log_2_ fold-change (log_2_FC) values were computed and log_2_FC values were then scaled by its standard deviation to derive log_2_FC-z scores^19,21^. Statistical significance was defined by the FDR threshold of 0.05 and an absolute log_2_FC-z cutoff of 2.

## Data Records

All data generated in this study are publicly available in the MassIVE repository (https://massive.ucsd.edu/) under accession number MSV000099982. The deposited dataset includes the following components:

The deposit includes the following components:

1. Raw Mass Spectrometry Files: All raw data files from the mass spectrometer.
2. Database Search Results: Output files from the JUMP software suite, including peptide and protein identification results.
3. Processed Quantification Matrix: A tab-delimited file containing quantified protein abundances. Rows correspond to proteins and columns correspond to individual samples. Matrix entries represent normalized TMT reporter ion intensities.
4. Metadata File: A metadata.txt file describing all sample-level attributes. This file links each SampleID in the quantification matrix to patient metadata and relevant technical variables, including:

- *SampleID*: Unique identifier matching the data matrix.
- *BatchID*: TMT batch identifier.
- *Patient_ID*: Anonymized patient identifier.
- *Age*: Age of the patient.
- *Sex*: Sex of the patient.
- *Race*: Self-reported race or ethnic background.
- *Risk*: Clinical risk stratum.
- *Diagnosis*: ALL types (for example, B-ALL or T-ALL).
- *Treatment*: Sample collection timepoint (pre- or post-treatment).

## Technical Validation

### Balanced and controlled quantitative proteomics design

We designed a quantitative proteomics experiment on cerebrospinal fluid (CSF) samples from pediatric acute lymphoblastic leukemia (ALL) patients, collected before and after treatment across ALL subtypes and risk strata (Fig. 1). The study was structured to enable comparisons across treatment stages, ALL types, and risk levels. Samples were prepared in 16 sample batches, each containing 15 patient CSF samples and one internal reference pooled sample. To prevent batch-related bias, a block randomization strategy was applied^22,23^, and all CSF samples were processed uniformly across 24 distinct batches.

**Figure 1:**
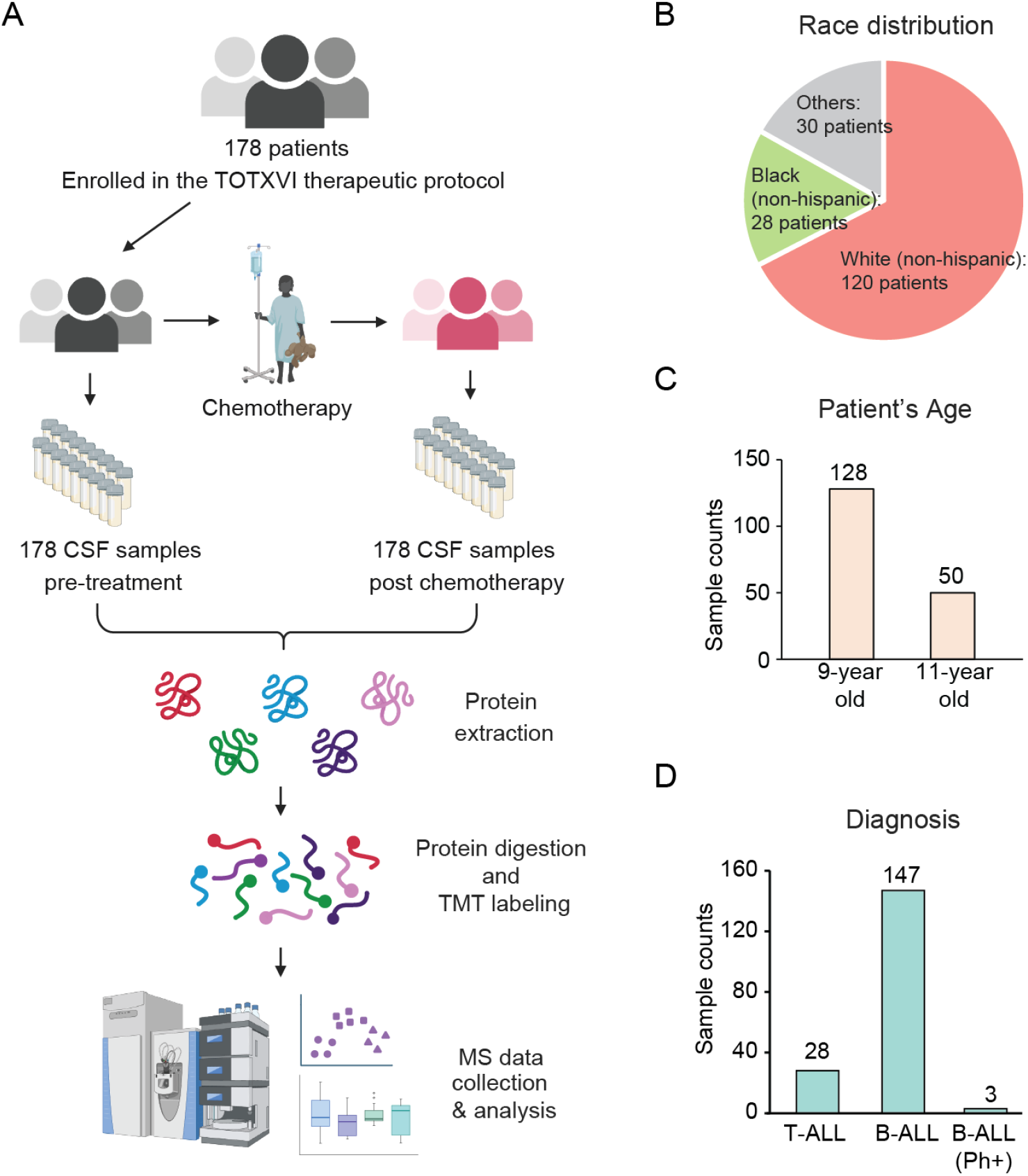
Overview of the TOTXVI Patient Cohort and CSF Proteomics Workflow. (A) CSF proteomics workflow. CSF samples were collected pre- and post-chemotherapy from 178 patients enrolled in the TOTXVI therapeutic protocol. Proteins were analyzed using TMTpro. (B) Race distribution of the patient cohort. Most patients were White (non-hispanic) (n=120), followed by Black (non-hispanic) (n=28), and Others (n=30). (C) Patient age distribution. The cohort predominantly consisted of 9-year-old patients (n=128) and 11-year-old patients (n=50). (D) Diagnosis distribution of the patient cohort. The diagnoses included T-ALL (n=28), B-ALL (n=147), and B-ALL (Ph+) (n=3).

Protein concentrations were measured using short SDS-PAGE gel against a BSA standard curve. Digestion efficiency was evaluated on a small aliquot of each sample; if LC-MS/MS analysis revealed a miscleavage rate above 15%, additional trypsin was added. TMT labeling efficiency was verified by comparing a small aliquot to an unlabeled control, ensuring the absence of unlabeled peptides before quenching the reaction. To correct quantification variations and achieve equal sample representation in the multiplexed pool, a multi-step normalization strategy was implemented. First, a “premix ratio test” was performed on a small pooled aliquot of all TMT-labeled samples. Reporter ion intensities from this test guided the preparation of a second, adjusted mixture, which was then used to generate the final bulk pool, ensuring a balanced contribution from each sample. Prior to analyzing the fractionated experimental samples, LC-MS/MS system performance was validated using standard rat brain proteome. Key parameters were monitored which include LC system pressure, MS signal intensity, mass accuracy, and peak width, confirming that the instrument optimal operation.

### Quality evaluation of mass spectrometry data

Our quantitative proteomics workflow produced a deep and uniform dataset. Across 24 TMT batches and 960 MS runs, we identified 9,356 proteins supported by 7,034,782 PSMs, corresponding to an average of 293,116 PSMs per batch. Applying a 50 percent data completeness threshold yielded >7,400 reliably quantified proteins. To ensure biological specificity and avoid inflation from highly similar gene products, we consolidated protein isoforms, resulting in a curated high-confidence set of 3,188 proteins. These proteins were highly consistent, with each of the 24 batches identifying around 3,000 proteins (Fig. 2A). Principal component analysis (PCA) revealed a thorough intermixing of all 24 batches, indicating the absence of significant batch-driven artifacts (Fig. 2B). The Log_2_ intensity distributions across all samples were highly uniform, confirming minimal technical variability and equal loading (Fig. 2C). Collectively, these metrics demonstrate that the dataset is high-quality, reproducible, and suitable for downstream biological analyses.

**Figure 2:**
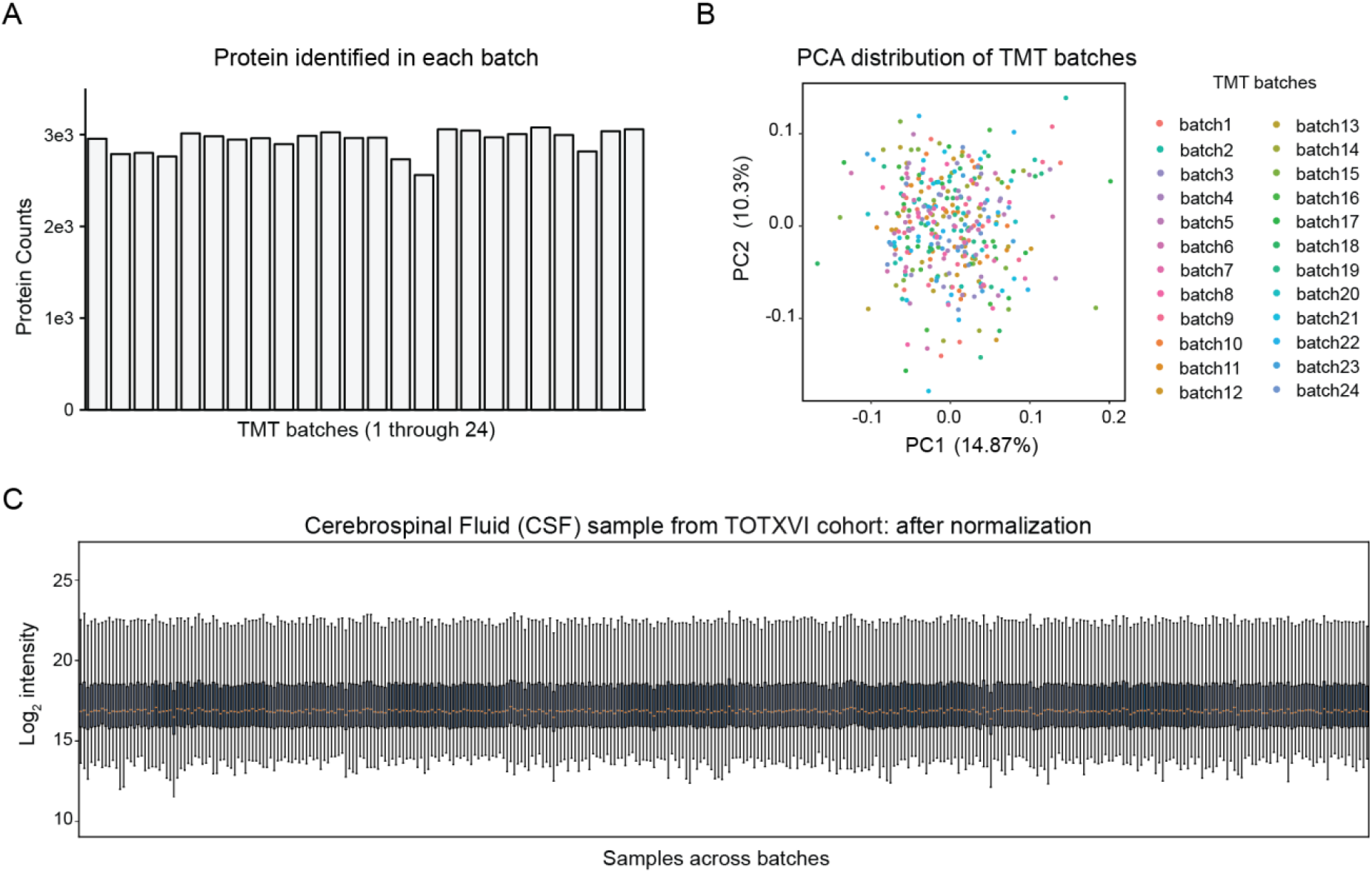
Quality Evaluation of Mass Spectrometry Data. (A) Protein identification consistency across TMT batches. Bar chart displaying the total number of proteins identified in each of the 24 TMT batches. (B) PCA distribution of TMT batches. A principal component analysis (PCA) plot showing all samples, colored by their respective TMT batch. (C) Log_2_ intensity distribution across the samples. Box plots illustrate the uniform alignment suggesting the equal sample loading and minimal technical variability.

To further ensure data integrity and mitigate confounding effects, standard quality control and processing steps were implemented (Fig. 3A). Coefficient of variation (CV) were calculated for the dataset which centered around 2-3% suggesting high technical precision (Fig. 3B). To ensure that downstream analyses were not confounded by non-biological factors, we computationally corrected for the effects of covariates including age, sex, and batch using a robust linear model. The success of this correction was validated by analyzing the p-value distributions for protein associations before and after correction (Fig. 3C and 3D). Finally, the dataset was assessed for abundance and dynamic range.

**Figure 3:**
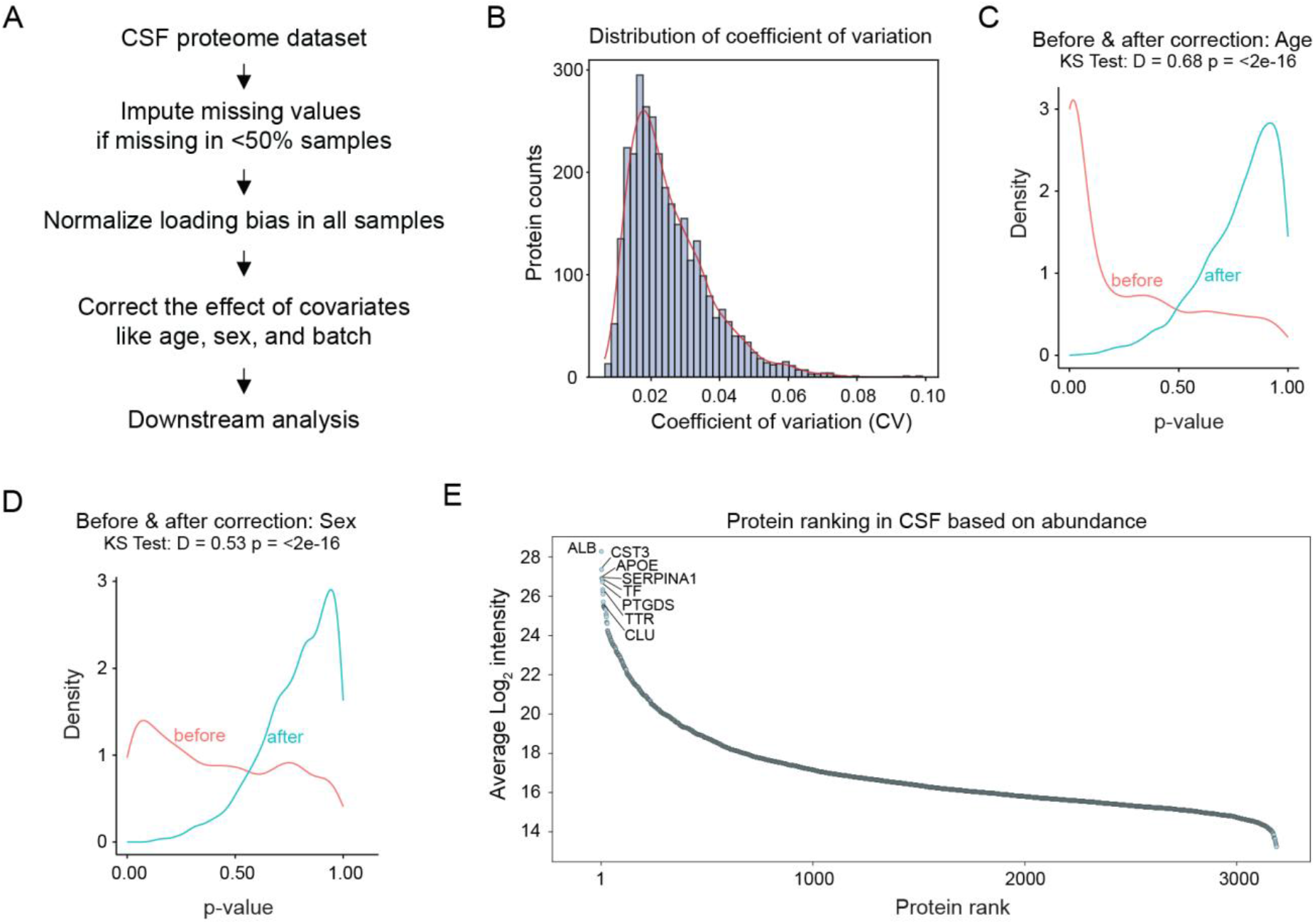
Data Processing and Proteome Characteristics. (A) Data processing workflow. A flowchart outlining the key computational steps applied to the CSF proteome dataset. (B) Distribution of coefficient of variation (CV). Histogram of the CVs for all quantified proteins. The distribution is tightly centered around 2-3%, indicating high technical precision and data reproducibility. (C-D) Validation of covariate correction. Density plots showing the p-value distributions for protein associations before (red) and after (blue) computational correction for (C) Age and (D) Sex. (E) Protein abundance ranking in CSF.

## Data Overview

### CSF proteomics overview

The final dataset of 3,188 proteins spanned a broad dynamic range typical of CSF, with abundances covering several orders of magnitude (Fig. 3E). A rank-abundance plot, ordering proteins from highest to lowest average log_2_ intensity, highlights well-known CSF proteins as top ranked. The most abundant protein in our dataset was Albumin (ALB), a well-known and highly abundant component of CSF^24^. Other highly ranked proteins, including Cystatin C (CST3), Apolipoprotein E (APOE), and Transthyretin (TTR), which are also characteristic high-abundance proteins found in CSF^25-27^. This profile confirms that the mass spectrometry analysis successfully captured the full breadth of the proteome, from high-abundance proteins down to low-abundance species, validating the depth of coverage.

### Protein differential abundant analysis

To identify chemotherapy-induced alterations in the CSF proteome, differentially abundant proteins (DAPs) were performed using a moderate paired t-test with a significance threshold of FDR < 0.05 and an absolute log_2_FC-z ≥2. This analysis detected 460 DAPs between post- and pre-treatment samples, comprising 222 upregulated and 238 downregulated proteins (Fig. 4A). Notably, the most strongly upregulated proteins included GPAT2 and several apolipoproteins (APOA1, APOA2, APOC1), while downregulated proteins were prominently represented by SELL, CRP, and ICAM1. These data clearly demonstrated the profound impact of chemotherapy on the CSF proteome. To incorporate these changes to the functional perspective, we performed pathway enrichment analysis on upregulated and downregulated DAPs. Subsequent pathway enrichment analysis (Fig. 4B) revealed that upregulated DAPs are significantly associated with cholesterol metabolism, complement and coagulation cascades, and platelet activation. Conversely, downregulated DAPs were primarily enriched in pathways related to nucleosome assembly and leukocyte cell-cell adhesion. Further validation of the top three upregulated proteins (GPAT2, APOC1, and APOA2) confirmed their consistent and significant increase post-treatment across all tested ALL subtypes, including T-ALL, B-ALL, and B-ALL(Ph+) (Fig. 4C), indicating a general therapeutic response.

**Figure 4:**
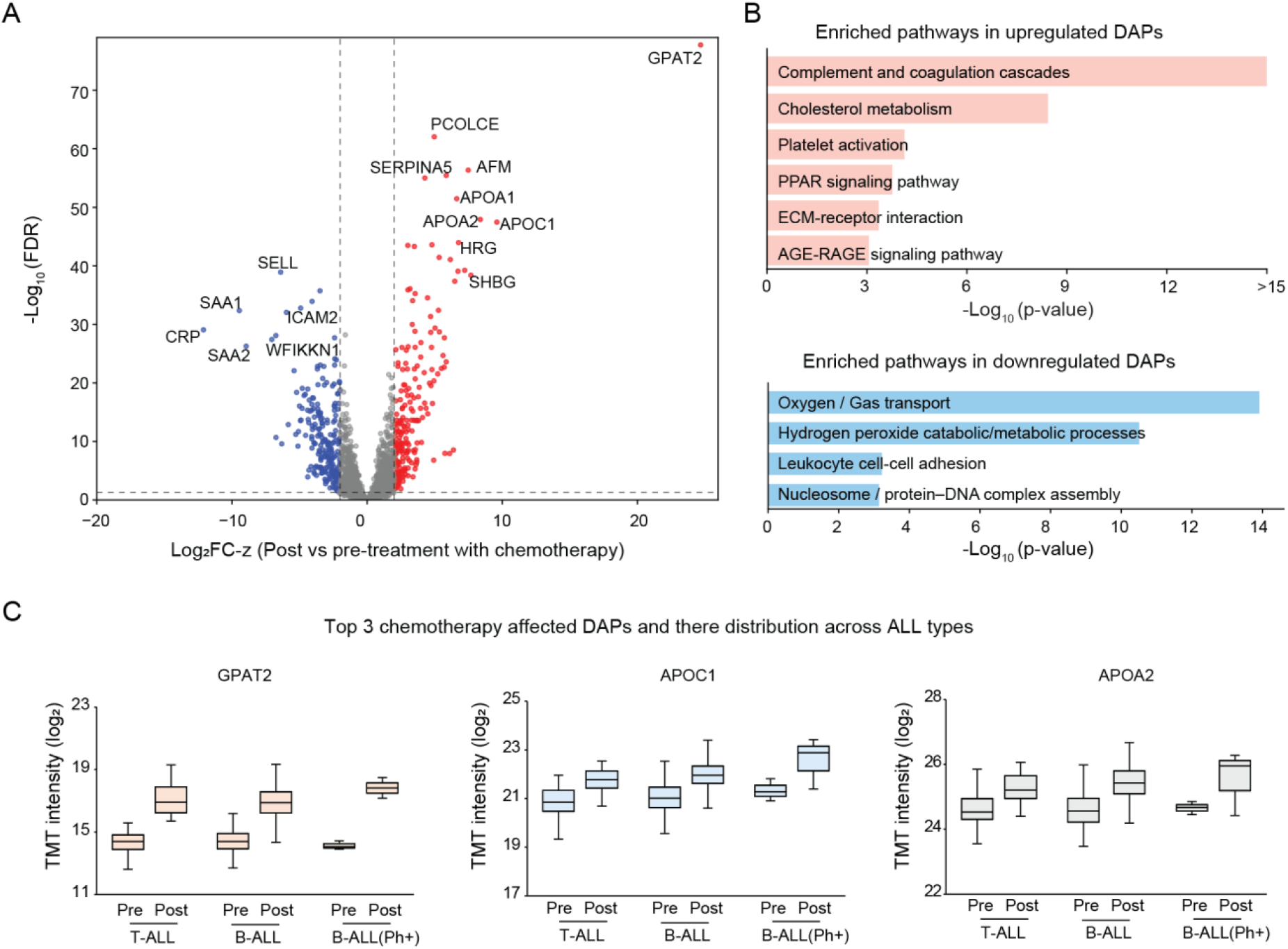
Identification of Chemotherapy-Affected Proteins. (A) Volcano plot of DAPs. A volcano plot displaying the DAPs between post-treatment and pre-treatment CSF samples. Red dots represent significantly upregulated DAPs (Log_2_FC-z > 2, FDR < 0.05), blue dots represent significantly downregulated DAPs (Log_2_FC-z < -2, FDR < 0.05), and grey dots indicate non-significant proteins. (B) Enriched pathways in upregulated and downregulated DAPs. The - Log10(p-value) for each process is indicated. (C) Distribution of top 3 chemotherapy-affected DAPs across ALL types. Box plots illustrate the TMT intensity (log_2_) of the three most significant chemotherapy-affected DAPs (GPAT2, APOC1, and APOA2) in pre- and post-treatment samples across different ALL types (T-ALL, B-ALL, B-ALL(Ph+)). Each plot shows the median and interquartile range, demonstrating their changes in abundance relative to chemotherapy and ALL subtypes.

## Usage Notes

Proteomics data generated from search engines tools require essential pre-processing steps before any downstream statistical analysis or biological interpretation. To support diverse analytical needs, the accompanying protein quantification table (Table S1-S4) includes several data tables, such as loading-normalized and log_2_-transformed TMT reporter intensities, an imputed dataset (in which missing values have been estimated if missing in <50% sample), and a covariate-corrected dataset (in which technical and biological effects, including age, sex, and batch, have been regressed out). Users should select the appropriate data set based on their analytical goals. For example, filtering out proteins with any missing values yields a high-confidence matrix composed exclusively of directly quantified measurements, but at the cost of reduced proteome depth. In contrast, imputed datasets retain broad proteome coverage but incorporate statistically estimated values. Covariate correction is strongly recommended, as unadjusted confounders can obscure biological signals; users may also apply alternative, widely used correction approaches such as ComBat^28^ or QC-RLSC^29^. Beyond age, sex, and batch effects, race may also be included as a covariate. Regardless of the chosen pre-processing strategy, users must ensure that all analytical decisions respect the original experimental design (for example, paired sampling), as these design features critically influence appropriate statistical tests and interpretation of results.

## Ethics statement

This study involves human participants. All patients were enrolled on the St. Jude Children’s Research Hospital institutional therapeutic protocol TOTXVI. The study was conducted in accordance with the Declaration of Helsinki and was approved by the St. Jude Institutional Review Board (IRB). Informed consent was obtained from the parents or legal guardians of all participants, with assent obtained from patients as appropriate for their age.

## Data Availability

The mass spectrometry proteomics data (raw files, search results, metadata, and other associated files) have been deposited in the ProteomeXchange Consortium via the MassIVE repository under the accession number MSV000099982. The dataset can be accessed through the MassIVE site (https://massive.ucsd.edu/) by searching the accession number, or directly at https://massive.ucsd.edu/ProteoSAFe/dataset.jsp?task=23d13497fed7420093b016fdd4bcdeee.

An FTP download option is also available at ftp://massive-ftp.ucsd.edu/v11/MSV000099982/.

## Code Availability

Proteomic data was processed using JUMP software suite that are publicly available (github.com/JUMPSuite).

## Author Contributions

M.N. and H.W. generated mass spectrometry data. H.K.S. performed data analysis and wrote the manuscript along with J.P. X.W., H.W., and J.P. assisted in the data analysis. K.K and J.P. conceived, supervised the project, and secured funding.

## Acknowledgements

We thank Dr. Xiaojun Sun for his assistance with sample processing. We are grateful to Dr. Justin Tanner for providing scientific feedback on the manuscript. We also thank the St. Jude Biorepository for providing the sample resources. Finally, we acknowledge all members of the Peng lab and the Center for Proteomics and Metabolomics at St. Jude Children’s Research Hospital for their invaluable technical support and insightful discussions.

